# TAP: Targeting and analysis pipeline for optimization and verification of coil placement in transcranial magnetic stimulation

**DOI:** 10.1101/2021.05.09.443339

**Authors:** Moritz Dannhauer, Ziping Huang, Lysianne Beynel, Eleanor Wood, Noreen Bukhari-Parlakturk, Angel V. Peterchev

## Abstract

**Objective:** Transcranial magnetic stimulation (TMS) can modulate brain function via an electric field (E-field) induced in a region of interest (ROI) in the brain. The ROI E-field can be computationally maximized and set to match a specific reference using individualized head models to find the optimal coil placement and stimulus intensity. However, the available software lacks many practical features for prospective planning of TMS interventions and retrospective evaluation of the experimental targeting accuracy.

**Approach:** The TMS targeting and analysis pipeline (TAP) software uses an MRI/fMRI-derived brain target to optimize coil placement considering experimental parameters such as the subject’s hair thickness and coil placement restrictions. The coil placement optimization is implemented in SimNIBS 3.2, for which an additional graphical user interface (TargetingNavigator) is provided to visualize and adjust procedural parameters. The coil optimization process also computes the E-field at the target, allowing the selection of the TMS device intensity setting to achieve a specific E-field strength. The optimized coil placement information is prepared for neuronavigation software, which supports targeting during the TMS procedure. The neuronavigation system can record the coil placement during the experiment, and these data can be processed in TAP to quantify the accuracy of the experimental TMS coil placement and induced E-field.

**Main results:** TAP was demonstrated in a study consisting of three repetitive TMS sessions in five subjects. TMS was delivered by an experienced operator under neuronavigation with the computationally optimized coil placement. Analysis of the experimental accuracy from the recorded neuronavigation data indicated coil location and orientation deviations up to about 2 mm and 2°, respectively, resulting in an 8% median decrease in the target E-field magnitude compared to the optimal placement.

**Significance:** TAP supports navigated TMS with a variety of features for rigorous and reproducible stimulation delivery, including planning and evaluation of coil placement and intensity selection for E-field-based dosing.

## 1. Introduction

Precise placement of non-invasive brain stimulation devices, such as transcranial magnetic (TMS), offers promising avenues for understanding basic (e.g., [1]) and higher (e.g., [2]) cognitive brain functions. In neuroscience experiments, TMS coil placement is often landmark-referenced and supported by neuronavigation technology [3,4], targeting a particular brain region of interest (ROI). These ROIs can be determined individually by identifying anatomical brain structures and their connectivity with MRI [4–6] or from the brain activity or functional connectivity obtained with fMRI or EEG [2,7–16].

Prior to a TMS session, individualized computational modeling of head tissues and the electric field (E-field) induced in the ROI [17] allows for the identification of an optimal TMS coil placement on the scalp [12,18]. We previously developed the fast auxiliary dipole method (ADM) for software-assisted TMS targeting to maximize E-field delivery to an ROI [18], and ADM is now part of the SimNIBS E-field simulation software package. However, it remains challenging to combine ADM with individual functional or anatomical targeting data, subsequent experimental application using neuronavigation, and visualization and evaluation of the experimental accuracy. Previous publications have proposed proof-of-principle fMRI-based TMS targeting pipelines [12,19]. However, these efforts have several limitations with regard to coordinate transformations across software packages, coil placement constraints and optimization, inclusion of hair thickness information, user interface for verification of ROI input information, and retrospective validation of TMS coil placement accuracy.

We present a TMS targeting and analysis pipeline (TAP) software that helps bridge the gaps between individual imaging data, SimNIBS, and neuronavigation. SimNIBS can robustly generate head tissue models [20] from MRI data. The ADM method in SimNIBS determines the optimal TMS coil position and orientation on each subject’s scalp to maximize the magnitude or a directional component of the ROI E-field [18]. In addition to SimNIBS’ basic functionality for adjustment and visualization of ROIs, the coil, and desired E-field parameters, TAP offers a graphical user interface (GUI), called ‘TargetingNavigator’.

TAP can be used for prospective targeting optimization before an experiment and/or retrospective targeting analysis after an experiment. The first step, in a prospective approach for TMS coil placement or E-field simulation, is to extract an ROI center from a volumetric target mask (i.e., a nifti file) for SimNIBS. In this approach, TAP generates specifically formatted ASCII text files readable by TMS neuronavigation software [21]. The presented implementation is for the Brainsight neuronavigation system, but TAP can be adapted to other systems as well. TAP can also control practical aspects of coil placement, such as the coil handle direction, which ideally should not block the TMS coil tracker or the subject’s sight, which may need adjustment to control the induced E-field direction in the cortex [22,23]. Retrospective analysis uses TMS coil placement data from an existing experiment recorded by neuronavigation software. In retrospective analysis, TAP can detect problematic coil distances from the scalp, such as a coil placement being too far from or entering the scalp surface due to inaccurate coil–scalp coregistration, or movement of the TMS coil during the stimulation; this analysis can incorporate measurements of the subjects’ hair thickness, if such are available.

In this paper, we describe TAP and illustrate its use in five TMS subjects who participated in a study of writer’s cramp dystonia.

## 2. Method

As shown in Figure 1A, TAP combines existing software packages, including SimNIBS v3.2 [20] for TMS-induced E-field simulations and Brainsight v2.5b2 [21] for neuronavigation, with custom MATLAB R2018a [24] code. The following sections describe how TAP was utilized in a complex study paradigm. Briefly, individual TMS target ROIs were derived from fMRI (Section 2.1), optimal coil placements were computed prospectively (Section 2.2) and applied in experimental TMS sessions (section 2.3), and the experimental coil placements were recorded and analyzed to assess the targeting accuracy (Section 2.4).

**Figure 1.**
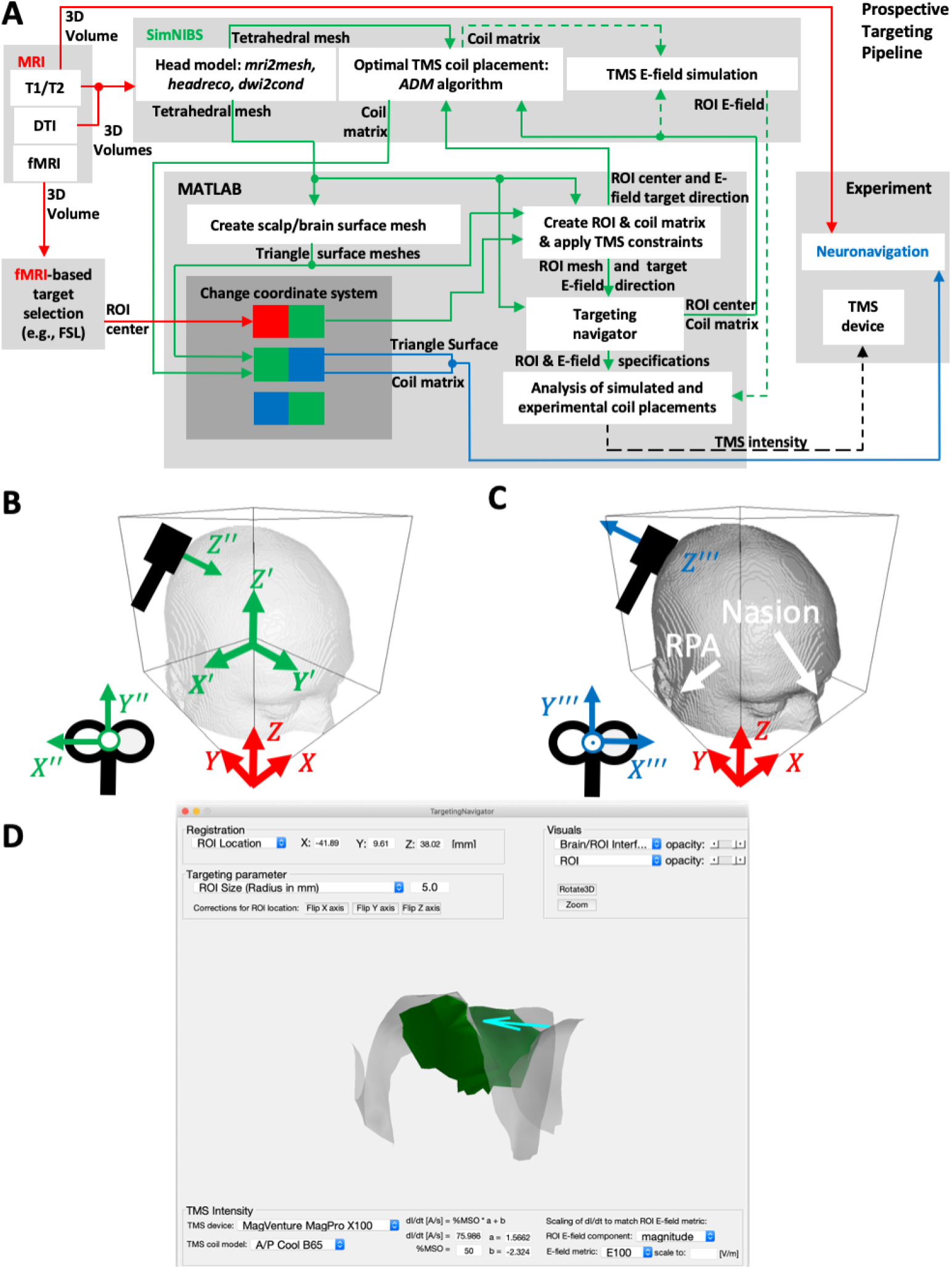
(A) Diagram of the TMS targeting and analysis pipeline (TAP) integrating different hardware and software components for prospective targeting. The dashed arrows indicate optional steps that are not required for prospective targeting. (B) SimNIBS software coordinate convention (green) with origin at the volumetric center, *X*’ and *Y*’ axes flipped with respect to the MRI RAI convention (X, Y, Z, in red), and a TMS coil coordinate system (*X*’”, *Y*’”, *Z*’”). (C) Neuronavigation (Brainsight) coordinate convention (blue) with pitch *X*’” = –*X*” and yaw *Z*’” = – *Z*” flipped compared to the SimNIBS convention. The nasion and right/left periauricular points (RPA/LPA) are typical registration points for neuronavigation. (D) A screenshot of the TargetingNavigator GUI: a MATLAB-based SimNIBS 3.2 add-on to adjust and visualize simulation parameters in prospective and retrospective TMS analysis.

### 2.1 Experimental participants, MRI, and target selection

#### 2.1.1 Participants⍰

The study [25] was open to all patients with writer’s cramp (WC) dystonia in their right hand who were not on any symptomatic treatments for more than three months and had no contraindications to MRI and TMS. Age-matched healthy volunteers (HV) who were right-hand dominant with no brain disorders were also recruited for the research study. Thirteen WC and 13 HV consented and completed the fMRI visit; of these, five subjects (four WC and one HV) consented and completed the three TMS visits as detailed below.

#### 2.1.2 fMRI research design⍰

*All subjects* completed a writing task in the MRI scanner during which they were presented with visual instructions to copy a sentence on an MRI-compatible digital tablet. The fMRI acquisition lasted for approximately 7.5 minutes and comprised twelve 20-second blocks of writing separated by periods of rest.

#### 2.1.3 Image acquisition and preprocessing

A total of 13 WC and 13 HV patients completed the structural MRI and functional MRI (fMRI) sequences. A 3 Tesla GE scanner was used to collect structural T1-weighted (echo-planar sequence: voxel size = 1 × 1 × 1.3 mm^3^, repetition time (TR) = 6.836 s; echo time (TE) = 2.976 s; field of view (FOV) = 25.6 mm^2^, bandwidth 41.67 Hz/pixel), T2-weighted (echo-planar sequence voxel size = 1 × 1 × 1.3 mm^3^, TR = 1267.0 ms, TE = 20 ms, flip angle 10°, FOV = 25.6 mm^2^, bandwidth 31.26 Hz/pixel), and diffusion-weighted scans (voxel size = 1 × 1 × 1.3 mm^3^, TR = 9000 ms, TE = 92.1 ms, FOV = 25.6 mm, bandwidth 250 Hz/pixel, matrix size 128 × 128, B-value = 2000 s/mm^2^, diffusion directions = 36). Functional echo-planar images were acquired while subjects performed the writing task using the following parameters: voxel size = 3.5 × 3.5 × 4.0 mm^3^, TR = 2 s, TE = 30 ms, flip angle = 90°, FOV = 22 cm, bandwidth = 250 Hz/Pixel, interleaved slice order with whole brain coverage). The Cigal software was used to align each subject’s movement to a reference frame to monitor movement in real time [26]. All fMRI images were preprocessed using fMRIPrep software after discarding the first five acquired brain volumes [27].

#### 2.1.4 TMS target selection

Preprocessed fMRI sequences were imported into the FSL software ([28], available at https://fsl.fmrib.ox.ac.uk/fsl/fslwiki) to perform general linear modeling (GLM) at the individual subject level (with spatial smoothing of 5 mm, cluster z-threshold of 2.3, and p-value of 0.05). The GLM model involved convolution of the writing blocks to the double-gamma hemodynamic response function to identify the regions of brain activation for each subject during the writing task compared to the rest period [26,28]. The z-stat brain activation maps of each subject’s writing task were then moved into a second-level group analysis using mixed effects FLAME1 modeling to generate the mean brain activation maps for 13 WC and 13 HV subjects during the writing task. The z-stat brain activation maps for each of the five subjects (four WC and one HV) were then visualized to develop customized ROIs for TMS stimulation. Specifically, each subject’s activation map was overlaid on the respective group activation map and a mask of the structural left premotor cortex or the primary sensory cortex as defined by the Harvard-Oxford MNI atlas [29]. ROI targets for TMS were defined by selecting two neighboring voxels, which were within the atlas-defined left premotor cortex or primary sensory cortex, had peak activation with z-statistic > 2.3, showed overlap between the individual and group activation maps, and were within 4 mm from the cortical surface. The 4 mm rule was used to ensure that the cortical ROI could be reached by the TMS E-field [3]. The individual ROIs were then reversed-transformed into individual native space using FLIRT linear registration with a sinc interpolation. The ROIs in the individual native MRI voxel space were then imported into the TAP workflow to identify the ROI center and transform to the computational model mesh (SimNIBS space) to determine the optimal TMS coil placements.

### 2.2 TMS targeting and analysis pipeline (TAP)

#### 2.2.1 Coordinate system transformation

Generally, coordinate transformations are needed to integrate all different software pieces (MRI, SimNIBS, and Brainsight) in the pipeline, mediated by custom MATLAB code. The coordinate transformations are defined as 4 × 4 matrices (denoted as *T_i,j_* with *i,j* = 1,…,4), which contain submatrices for rotation (*T_i,j_* with *i,j* = 1,2,3) defining axis rotations (with ||*T*_*,*j*_|| = 1 for *j* = 1,2,3) and translation (*T_i,j_* with *i* = 1,2,3; *j* = 4 in units of mm; *T*_4,4_ = 1), adding a coordinate offset that moves the origin of the current coordinate system to the specified desired one. The matrix *T_i,j_* is used to map locations such as the ROI and the TMS coil center, coordinate space rotations (i.e., colored arrows or letters in Figure 1A-C), and coil coordinate transformations from neuronavigation [21] or canonical coil space into SimNIBS modeling space [20], denoted as ‘Coil matrix’ in Figure 1A. The coordinate system transformations from MRI or Brainsight to SimNIBS space are performed based on mapping coordinates (expressed as world coordinates) relative to the center of the image volume of the respective MRI file created with SimNIBS (*SubjectName_T1fs_conform.nii.gz*) using either the mri2mesh or headreco pipeline. In more detail, *SubjectName_T1fs_conform.nii.gz* is present in the RPI coordinate/voxel-order system (except for the headreco option non-conform where it keeps the coordinate system of the input MRI), where the origin is at the RPI corner of the voxel space with positive voxels *X, Y* and *Z* axes, respectively. A 4 × 4 transformation matrix or quaternion transformation is stored in either MRI or ROI-defining image file (*nifti* file, extension: *nii.gz*) with additional information that is used by default to transform voxel coordinates to world coordinates. The nifti-header information can be modified by a variety of image-processing software, and some may transform any image to one specific voxel-order/coordinate system (e.g., to the RAI in Seg3D, [30]), creating the appearance of being at a proper location. To ensure the correctness of the transformed ROI coordinate, TAP computes the shortest Euclidean distance to the next segmented gray matter volume in SimNIBS MRI-space (i.e., ‘*SubjectName_T1fs_conform.nii.gzf*’) using ‘*gm _fromMesh.nii.gz*’, SimNIBS v3 or ‘*gm_only.nii.gz*’ for SimNIBS v2 mri2mesh runs. If this distance is larger than the voxel size, then the ROI and gray matter do not overlap. In this problematic case, which is most likely due to voxel-order/ coordinate transformation misalignment, TAP can be configured to systematically sign-flip the axes/origin to find a world coordinate transformation with minimal distance or do that manually by starting the TargetingNavigator for inspection and adjustment.

Once the ROI voxels are in SimNIBS MRI-space (i.e., in *SubjectName_T1fs_conform.nii.gz*) as world coordinates (*X,Y,Z*), they need to be converted to SimNIBS model space (*X*’, *Y*’, *Z*’) by representing the coordinates relative to the image center and flip axis (*X*’ = –*X,Y*’ = –*Y,Z*’ = *Z*, see Figure 1B). The physical orientation of the TMS coil is defined in the SimNIBS space (denoted *X*”, *Y*”, *Z*”) and Brainsight space (denoted as *X*’”, *Y*’”, *Z*’”), while flipping by 180° the pitch (*X*’” = – *X*”) and yaw coil orientation axes (i.e., *Z*’” = – *Z*”). Coil orientations are visualized by black-colored coil representations in Figure 1B and 1C, respectively. For biphasic TMS pulses, TMS is most effective when the direction of the middle phase of the induced E-field points to the wall of the cortical sulci [2,23,31,32]. As described in section ‘Preparation and launch of SimNIBS’, TAP estimates this direction based on the gyral/sulcal transition zone in the ROI (i.e., through cortical curvature information in SimNIBS mri2mesh; for more details, see below) to find a representative location on the wall for which it computes a smoothed inward-pointing normal vector. This vector is then used as the ROI directional E-field constraint in the ADM algorithm to determine the optimal TMS coil setup. After ADM has finished, the optimal coil orientation is prepared for Brainsight considering the following experimental constraints: The *Z*’” axis is checked for pointing in the scalp-outward direction (indicated visually by a large green cylinder cone in Brainsight). Further, for TMS devices that allow electronic reversal of the coil current direction, there is the option to flip the coil orientation and current direction for most practical experimental coil placements. It is advantageous to maintain the orientation of the TMS coil handle, including any attached cable, pointing away from the subject’s face, and TMS tracking headgear. To achieve this, TAP projects the optimized TMS coil orientation (*Y*”) onto the anterior-posterior *W*’ – *Z*’) plane and then computes an angle α with the *X*’ axis (equal to the zero-degree axis and positive angles for clockwise rotation). If 90° < α < 270°, then the coil orientation is flipped, and the necessity for coil current reversal is highlighted in the prepared Brainsight files.

#### 2.2.2 Preparation and launch of SimNIBS

TMS coil placement optimization can be launched with TAP by executing the *run_simnibs(s)* command from the MATLAB prompt (or within a script as in TAP) providing a MATLAB struct object *s* with different data fields, which should be initialized with the default struct *opt_struct*(‘*TMSoptimize*’). If s has the data field entry *s.target_direction*, ADM or the direct solver in SimNIBS will find the coil placement that maximizes the average E-field induced in the ROI along the vector *s.target_direction*, or otherwise the overall magnitude, as specified. The ROI center location in the SimNIBS space is assigned to the data field. The ROI size (*s.target_size*) considers all tetrahedral gray matter elements that fall within this radius (from the point specified in *s.target*) as Euclidian distance in three dimensions. For our five TMS subjects, an ROI size value between 3 and 4 mm was chosen to ensure that the ROI covered a significant part of the sulcal wall, but not any part of a neighboring gyrus controlled for by visual inspection in TargetingNavigator. The ROI target direction was determined using the following main steps:

1. project the ROI center onto the smoothed brain surface;
2. project location from (1) to the closest scalp surface mesh point and compute its nodal normal (averaging the neighboring triangle normal vectors) representing the coil plane.
3. find the sulcal wall location/trajectory (convex-concave transition of sulcus and gyrus):
  (3.1) mri2mesh: use location from (1) and Freesurfer’s surface curvature information (files: *lh.sulc, rh.sulc*; value range: between −2 and 2) to find the nearest surface mesh point that is numerically closest to zero.
  (3.2) headreco: use location from (1) and its scalp projection to form a trajectory vector.
4. compute an outwards-pointing nodal normal by averaging the triangle normal vectors for the brain surface nodes.
5. find sulcal wall surface normal and adjust for scalp coil plane:
  (5.1) mri2mesh: determine the nodal normal (4) for location (3.1).
  (5.2) headreco: move up/down the trajectory vector to find a location for which its nodal normal is mostly parallel to the tangential plane at the closest scalp node (2).

The sulcal wall normal vector found in (5) for mri2mesh or headreco, respectively, is then adjusted by projection onto the tangential plane (using the closest scalp node normal vector) to create the desired ROI target direction vector, because the TMS-induced E-field is mostly tangential to the scalp surface.

For robust estimation of the scalp-tangential plane, we added a data field to the s structure (available in SimNIBS 3.2.5), namely *s.scalp_normals_smoothing_steps* (TAP default is 20), to adjust the coil tangentiality to the scalp and visualize it in TargetingNavigator.

SimNIBS allows constraining the scalp search space for possible coil center positions and orientations, which TAP sets to the default values of *s.search_radius* = 25 mm, *s.spatial_resolution* = 1 mm, and *s.search_angle* = 1°. For a typical SimNIBS head model and the default values, the optimal coil placement search takes approximately 2 min on a regular laptop (Intel Core i5 1.6 GHz, 8 GB RAM) running ADM.

#### 2.2.3 Prospective TMS dosing

For the five TMS subjects, we used mri2mesh/dwi2cond (SimNIBS 2.0.1) to generate a five-tissue head model from the subject’s T1-weighted MRI data set, and assigned to the resulting tetrahedral mesh elements isotropic conductivities for the scalp (0.465 S/m), skull (0.01 S/m), and cerebrospinal fluid (1.654 S/m), as well as anisotropic conductivities for the gray and white brain matter. The anisotropic conductivities were derived from the subject’s co-registered diffusion-weighted MRI, where the diffusion tensors are volume-normalized so that the geometric mean of the conductivity tensor eigenvalues equals literature-based values of 0.275 S/m and 0.126 S/m for gray and white matter, respectively.

The subject’s hair thickness is often unknown before the TMS session [2], but can be measured at the beginning of the session. Therefore, multiple runs of SimNIBS’s ADM are required for a range of hair thickness values. The default setting in TAP iterates through a reasonable range of hair thicknesses from 0 to 7.5 mm in steps of 0.5 mm. The optimal coil setup was chosen for TMS targeting (i.e., as loaded into Brainsight) while a TMS intensity related to the subject’s resting motor threshold was employed as described in Section 2.3.

The TMS pulse intensity, specified as the coil current rate of change (*dI/dt*), can be set as a parameter directly in SimNIBS (default: *s.didt* = 10^6^ A/s = 1 A/μs) or through the TargetingNavigator interface. TAP also offers the option to determine the required *dI/dt* to reach a desired E-field strength in the ROI [2,34,35], which is an important feature for ROI-based dosing of TMS. The desired directional or magnitude E-field strength in the ROI can be specified in TAP by one of several metrics: the maximal value as well as statistically more robust metrics, such as the 50^th^, 75^th^, 90^th^, and 99^th^ percentiles, to avoid numerical E-field outliers. Our previously proposed E20, E50, and E100 metrics, which define the minimum E-field strength within the 20, 50, and 100 voxels with the strongest E-field in the ROI [2,36]. The computed *dI/dt* can be entered into the TMS device, typically as a percentage of the maximum stimulator output (%MSO), based on the approximately linear relationship between *dI/dt* and %MSO [2,36]. This relationship can generally be described by the formula

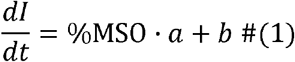

where coefficients *a* and *b* can be fitted to empirical data obtained from TMS devices that provide both %MSO and *dI/dt* [A/μs] readout. In this case, *dI/dt* should be measured across the full range of intensity for a specific coil, for example, with 5% MSO steps [2,36]. For devices that do not display *dI/dt,* the most practical approach is to set *b* = 0, and compute

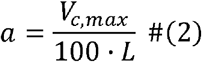

from the TMS device maximum capacitor voltage, **V**_C,max_, corresponding to 100 %MSO, and the specific coil inductance *L*, both of which can be obtained from the manufacturer. For example, *V_C,max_* is 1,800 V, 1,670 V, and 2,800 V for the standard MagVenture MagPro, Magstim Rapid^2^, and Magstim 200^2^ devices, respectively. The TargetingNavigator interface incorporates the ability to solve equation (1) for either *dI/dt* or %MSO with specified *a* and *b*, and provides values for these coefficients for some common TMS device and coil combinations.

#### 2.2.4 Neuronavigation files

TAP generates different types of files that can be visualized in Brainsight:

1. scalp and brain surface of the SimNIBS head model converted to MRI/Brainsight space,
2. SimNIBS/Freesurfer preprocessed T1-weighted MRI data set,
3. ROI/scalp coil center converted to MRI/Brainsight space, and
4. optimal coil placement (fully defined by a 4 × 4 matrix) was converted to the Brainsight space and saved in a text file.

For retrospective analysis, the experimenter saved the coil placement data as text files (i.e., for Brainsight software) during the TMS session, which can then be read into MATLAB and visualized in TargetingNavigator.

### 2.3 Experimental TMS sessions

The optimal TMS coil placement information of the five subjects who received TMS was imported as a project file (comprising PMC or PSC cortical target, T1, scalp/brain surface, and hair thickness estimations) into the Brainsight software for the TMS sessions. All TMS sessions were performed in a double-blind, sham-controlled, crossover research design. Three TMS sessions per subject were performed in a randomized order: active TMS at the premotor cortex, sham TMS at the premotor cortex, and active TMS at the primary sensory cortex, with each session one week apart to allow washout of the rTMS neuromodulatory effects. Each session was divided into four blocks, each 5 min apart. During each block, subjects received 25 trains of 3.9 s, 10 Hz repetitive TMS with an intertrain interval of 10 s at 90% resting motor threshold with biphasic stimulation for a total of 1,000 and 4,000 pulses per block and session, respectively. TMS was delivered using a MagPro X100 stimulator with an A/P Cool-B65 coil (MagVenture, Alpharetta, GA, USA). A neuronavigation system (Brainsight, Rogue Research, Montreal, Canada; [21]) was used to coregister the T1-MRI and the fMRI-based TMS target developed with TAP to the subject’s scalp. The location of the subject’s hot spot over the left motor cortex that induced the strongest motor evoked potentials in the contralateral first dorsal interosseus (FDI) was determined and marked in Brainsight. The resting motor threshold (rMT, expressed as %MSO) was then determined using a maximum-likelihood-based estimation procedure (see Table 1) [37]. The subject’s hair thickness over the TMS target area was measured (see Table 1) [2] and the closest hair thickness value in the reference table provided by TAP was selected. The TMS coil position and orientation values corresponding to the selected hair thickness were then used for each TMS session. During sham stimulation, the A/P coil was flipped, and subjects received similar auditory stimulation and scalp sensation through two surface electrodes placed on the subject’s scalp around the TMS target, producing only approximately 7.7% of the active E-field stimulation in the brain [38]. For all TMS sessions, the experimenter positioned the coil tangentially to the scalp surface with its handle directed posteriorly with a goal angle deviation < 5°. The coil placement data were saved by Brainsight every 500 ms (every fifth pulse) during the rTMS trains, totaling 200 recorded TMS coil setups per block, and exported as text files. Recording the coil position at this reduced rate compared to the pulse train rate was necessary for the neuronavigation system (Brainsight) to be able to save all data points correctly. No adverse events due to stimulation have been reported.

**Table 1.** rTMS intensity setting, equal to 90% of the individual resting motor threshold (rMT), and hair thickness measurement (in mm) of the five study subjects for the three TMS sessions (listed by target and TMS condition, not in chronological order).

| Subject | rTMS intensity setting (% MSO) |  |  | Hair thickness (mm) |  |  |
| --- | --- | --- | --- | --- | --- | --- |
|  | Session target & TMS condition |  |  | Session target & TMS condition |  |  |
|  | PMC Sham | PMC Active | PSC Active | PMC Sham | PMC Active | PSC Active |
| S1 | 52 | 53 | 47 | 0.5 | 0.5 | 0.5 |
| S2 | 76 | 69 | 64 | 2.5 | 2.0 | 2.5 |
| S3 | 39 | 41 | 38 | 1.5 | 1.5 | 2.5 |
| S4 | 46 | 50 | 47 | 2.0 | 2.5 | 2.0 |
| S5 | 55 | 58 | 45 | 2.0 | 2.0 | 3.5 |

### 2.4 Retrospective Analysis

The Brainsight neuronavigation system can digitize the coil position and orientation for each TMS pulse. TAP reads in (*read_brainsight_file*) the Brainsight-space exported text files and converts each line, corresponding to one digitized coil placement defined by a 4 × 4 transfer matrix, to SimNIBS modeling space (*convert_coord_brainsight_2_simnibs*). To avoid possible errors in the coil-scalp co-registration resulting in the coil center being inside the scalp, it is recommended to use Brainsight’s “Snap to Scalp” option when exporting [21]. TAP then extrudes these snapped coil centers outwards along the scalp normal to account for the measured hair thickness. If hair thickness measurement is unavailable, TAP results in a value of 2 mm [2]. The distance between the hair-thickness-extruded coil center and the original experimentally recorded coil center without snapping and hair-thickness extrusion was determined along the recorded coil normal direction (‘normal coil deviation’ in mm). The hair-thickness-extruded coil center is then compared against the computationally optimal coil center on the scalp’s tangential plane (‘tangential coil deviation’ in mm). TAP uses the computationally optimal coil placement to check if the rotational part of the experimental coil placement matrices aligns, and sign-flips the components for minimal angular discrepancy. The scalp normal direction was used to assess the deviation of the coil plane from the scalp-tangential plane (‘normal coil deviation’ in angular degree (°) equal to the arccosine of the dot product of the scalp and coil normal). Finally, the experimental coil placement is projected onto the scalp-tangential plane and the difference between the experimental and optimal coil orientation is computed (‘tangential coil deviation’ in angular degree (°)). When a targeted optimal coil placement is not available, the TAP can still analyze the experimental TMS coil placements recorded with Brainsight.

As a second step, TAP evaluates the difference in induced E-field between the optimal (if one is provided) and all experimentally recorded coil setups. Because one E-field simulation requires a computation time of several minutes, TAP avoids a very long total simulation time by choosing a median representative of the coil placements within an experimental block to simulate the Efield in the entire head model and then extracts the ROI E-field (using SimNIBS ‘target.msh’). The median coil representation is chosen by evaluating the medians for the *X*’”, *Y*’” and *Z*’” coordinates separately and picking the recorded coil placement with the smallest Euclidean distance from these median coordinates. For the E-field simulations, the value of *dI/dt* is computed with Equation (1) from the individual rMT in units of %MSO, and entered into the *s.didt* field of the MATLAB structure inputted to SimNIBS. For the TMS coil and device in this study, coefficient values of *a* = 1.5662 and *b* = −2.3237 were used in Equation (1) based on prior calibration data [2]. Finally, TAP computes the deviation of magnitude and angle of the E-field vector in each tetrahedral ROI element between the simulations for the median experimental coil placement and the optimal coil placement.

## 3. Results

Figure 2B-G quantifies the deviation between the computationally optimized coil placement that was set as a target during the TMS sessions and the actual placements recorded by the neuronavigation system. Across all subjects and sessions, the median coil position and orientation deviations were −1.55 mm and 1.28° relative to the scalp normal (Figure 2B,D) and 2.18 mm and −0.29° in the scalp tangential plane (Figure 2C,E), respectively. The corresponding median E-field magnitude and direction deviations were 7.69% and 0.65° (Figure 2F,G), respectively, even though some of the individual medians were significantly larger.

**Figure 2.**
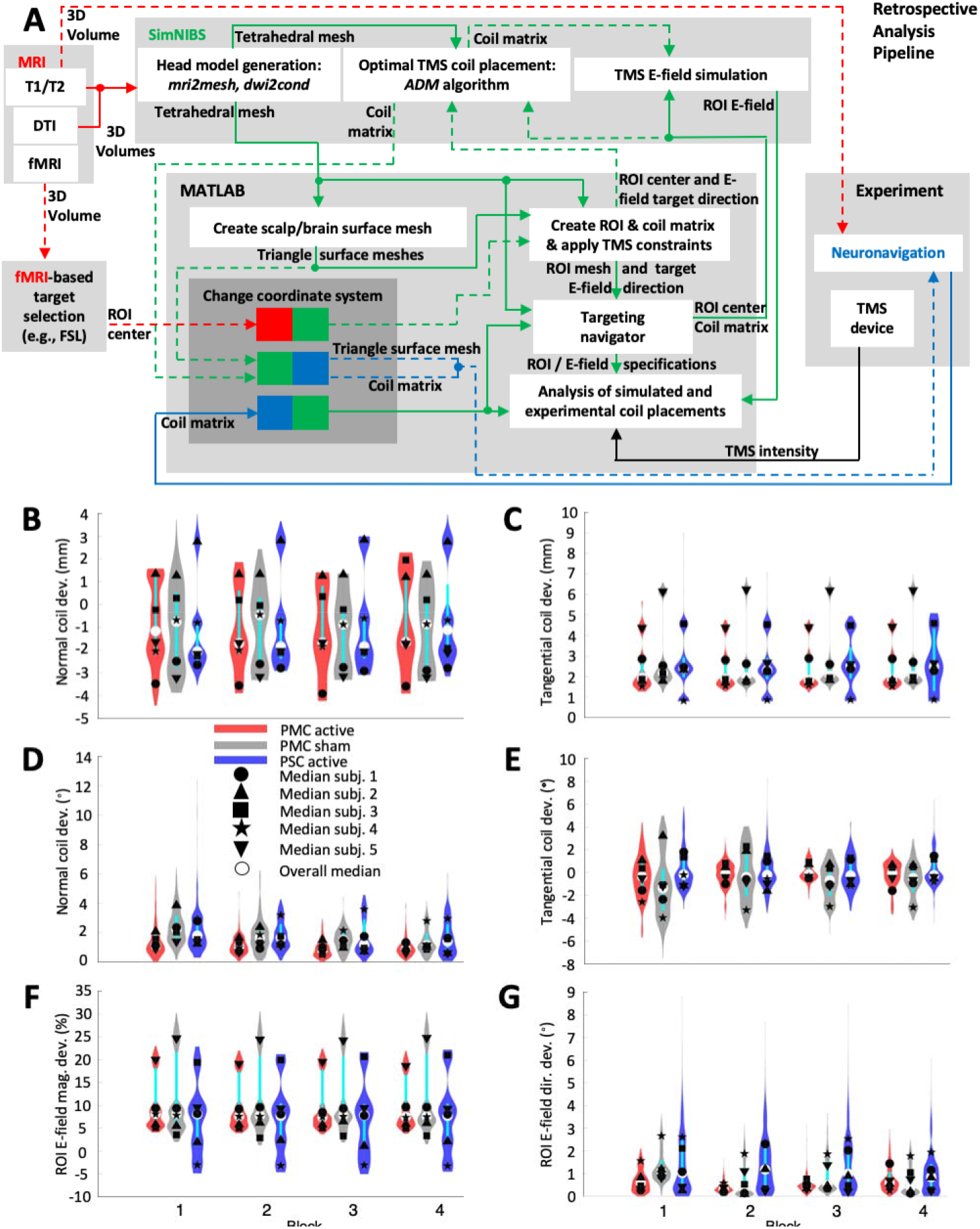
(A) Diagram of TAP as used for retrospective dosing analysis based on TMS coil scalp placement data recorded via neuronavigation (Brainsight). Dashed arrows indicate optional steps that are not required for retrospective TMS analysis. (B)–(G) Violin plots and medians of the differences in coil placement and induced E-field between prospectively-optimized and neuronavigation-recorded targeting in 5 subjects for 3 rTMS sessions of 4 blocks each. Deviation distance of (B) the coil center along the coil normal (yaw axis) direction (positive/negative values represent coil surface above/below the scalp surface) and (C) the coil center in the plane tangential to the scalp. Angular deviation of (D) the coil normal (yaw axis) and (E) the coil orientation about its yaw axis with clockwise as the positive direction. (F) Magnitude and (G) angular deviation of the E-field vectors for each finite element in the ROI. The violin plots show the distribution of the data, with thin black and thicker cyan vertical lines representing the 95% confidence interval and the interquartile range, respectively [33]. The plots exclude samples that have normal or tangential deviation distance of more than ± 10 mm, which were considered outliers that likely occurred due to brief disruptions in the coil tracking and comprised only 0.069% of the data.

These values are comparable to the neuronavigation error and deviation estimates reported in the literature. TMS neuronavigation systems are known to have a coil placement accuracy of 5-6 mm on average [39,40]. Excluding errors from the registration and shifts of the head tracker relative to the head, the median tangential deviation of the coil position and orientation of 1.34 mm and 3.48°, respectively, was reported for a neuronavigated robotic coil holder [41]. Moreover, an inter-session position error of approximately 2.5 mm was reported for neuronavigated manual coil placement [42].

## 4. Discussion

The novel targeting and analysis pipeline (TAP) integrates the SimNIBS E-field simulation package with the Brainsight neuronavigation system using custom MATLAB code to improve the practicality of precise brain targeting in TMS experiments using E-field simulations. The proposed capabilities of TAP software go beyond what has been previously published (e.g., [12]). TAP enables prospective and retrospective TMS targeting analysis based on E-field simulations coupled with individual imaging and neuronavigation data. Prospective targeting computations include the optimization of the TMS coil placement as well as the selection of the TMS pulse intensity to deliver a specific E-field strength at the target ROI. A retrospective analysis of the experimental coil placement can quantify the overall accuracy of applying TMS. For example, retrospective coil placement analysis can be used to remove problematic TMS sessions, blocks, or pulses, perform statistical analyses to explain experimental outcomes, and draw more accurate conclusions from hypothesis-driven TMS research. We demonstrated the use of TAP in an example study in which TMS coil positioning deviations did not exceed the error levels of neuronavigation systems. This, of course, depends on the precision of coil handling across experimental blocks, sessions, and subjects, and should be evaluated in each study. Moreover, the coil positioning deviations are detected only relative to the target placement set in the neuronavigation coordinates. The absolute precision relative to the actual brain target also depends on the accuracy of the neuronavigation registration. Therefore, the initial registration must be accurate, and it may be prudent to verify the registration of head landmarks and recorded scalp coil placements at the end and potentially within an experimental session [43].

TAP is an extension of SimNIBS, which is freely available. It is our hope that TAP will enable more researchers to utilize SimNIBS E-field simulation capabilities as part of rigorous TMS targeting and analysis methods. TAP also facilitates readout from the Brainsight neuronavigation system to uncover potential issues regarding coil placement, including coil tracking and/or head registration. Presently, the TAP code supports only Brainsight, but we aim to extend compatibility with other neuronavigation systems in the future. Finally, we welcome input and contributions to the TAP code by users through the contact on the TAP GitHub page linked in the Data & Code Availability section.

## 5. Conclusion

TAP facilitates prospective planning and retrospective analysis of TMS coil placement and E-field delivery to individual brain targets. This functionality enables both the selection of the TMS coil placement and intensity to standardize E-field exposure as well as the quantification of variability in the coil placement and E-field dose across different subjects and experimental sessions. We demonstrated the use of TAP in an example study in which TMS coil positioning deviations did not exceed the error levels of neuronavigation systems. TAP can be a useful tool for TMS practitioners to precisely plan dosing and analyze experiments or interventions to ensure rigorous and reproducible stimulation delivery.

## Acknowledgements

This work was supported by grants from the National Institutes of Health (RF1MH114268, RF1MH114253, U01AG050618, KL2TR002554), Dystonia Medical Research Foundation, Doris Duke Charitable Foundation, and Dystonia Coalition. The content is solely the responsibility of the authors and does not necessarily represent the official views of the funding agencies.

A. V. Peterchev has received research funding, travel support, patent royalties, consulting fees, equipment loans, hardware donations, and/or patent application support related to magnetic stimulation from Rogue Research, Tal Medical/Neurex, Magstim, MagVenture, Neuronetics, BTL Industries, and Advise Connect Inspire. The other authors report no relevant disclosures.

## Data and Code Availibilty

The software described herein as well as resulting data to recreate all figures are available on https://github.com/moritzdannhauer/TAP.git.

## Ethics Approval, Patient Consent, and Clinical Trial Registration

The human subjects participating in the fMRI and TMS experiments were part of a multi-session study of dystonia, including structural and functional magnetic resonance imaging and behavioral testing. The study was approved by the Duke University Health System Institutional Review Board (#0094131), ensuring ethical guidelines were followed, and all subjects provided written informed consent to participate. The study was not registered as a clinical trial because it was a small pilot feasibility study; moreover, this paper focuses on methodological aspects of TMS targeting and does not report any clinical results.

## References

[1] Jung J, Bungert A, Bowtell R and Jackson S R 2020 Modulating brain networks with transcranial magnetic stimulation over the primary motor cortex: a concurrent TMS/fMRI study Front. Hum. Neurosci. 14 31

[2] Beynel L, Davis S W, Crowell C A, Dannhauer M, Lim W, Palmer H, Hilbig S A, Brito A, Hile C, Luber B, Lisanby S H, Peterchev A V, Cabeza R and Appelbaum L G 2020 Site-specific effects of online rTMS during a working memory task in healthy older adults Brain Sci. 10 255

[3] Aberra A S, Wang B, Grill W M and Peterchev A V 2020 Simulation of transcranial magnetic stimulation in head model with morphologically-realistic cortical neurons Brain Stimulat. 13 175–89

[4] Rusjan P M, Barr M S, Farzan F, Arenovich T, Maller J J, Fitzgerald P B and Daskalakis Z J 2010 Optimal transcranial magnetic stimulation coil placement for targeting the dorsolateral prefrontal cortex using novel magnetic resonance image-guided neuronavigation Hum. Brain Mapp. 31 1643–52

[5] Nummenmaa A, McNab J A, Savadjiev P, Okada Y, Hämäläinen M S, Wang R, Wald L L, Pascual-Leone A, Wedeen V J and Raij T 2014 Targeting of white matter tracts with transcranial magnetic stimulation Brain Stimulat. 7 80–4

[6] Silva L M, Silva K M S, Lira-Bandeira W G, Costa-Ribeiro A C and Araújo-Neto S A 2021 Localizing the primary motor cortex of the hand by the 10-5 and 10-20 systems for neurostimulation: an MRI study Clin. EEG Neurosci. 52 427–35

[7] Wang J X, Rogers L M, Gross E Z, Ryals A J, Dokucu M E, Brandstatt K L, Hermiller M S and Voss J L 2014 Targeted enhancement of cortical-hippocampal brain networks and associative memory Science 345 1054–7

[8] Bergmann T O, Karabanov A, Hartwigsen G, Thielscher A and Siebner H R 2016 Combining non-invasive transcranial brain stimulation with neuroimaging and electrophysiology: current approaches and future perspectives Neuroimage 140 4–19

[9] Zrenner C, Belardinelli P, Müller-Dahlhaus F and Ziemann U 2016 Closed-loop neuroscience and non-invasive brain stimulation: a tale of two loops Front. Cell. Neurosci. 10 92

[10] Thut G, Bergmann T O, Fröhlich F, Soekadar S R, Brittain J-S, Valero-Cabré A, Sack A T, Miniussi C, Antal A and Siebner H R 2017 Guiding transcranial brain stimulation by EEG/MEG to interact with ongoing brain activity and associated functions: a position paper Clin. Neurophysiol. 128 843–57

[11] Curtin A, Tong S, Sun J, Wang J, Onaral B and Ayaz H 2019 A systematic review of integrated functional near-infrared spectroscopy (fNIRS) and transcranial magnetic stimulation (TMS) studies Front. Neurosci. 13 84

[12] Balderston N L, Roberts C, Beydler E M, Deng Z-D, Radman T, Luber B, Lisanby S H, Ernst M and Grillon C 2020 A generalized workflow for conducting electric field-optimized, fMRI-guided, transcranial magnetic stimulation Nat. Protoc. 15 3595–614

[13] Cash R F, Weigand A, Zalesky A, Siddiqi S H, Downar J, Fitzgerald P B and Fox M D 2020 Using brain imaging to improve spatial targeting of transcranial magnetic stimulation for depression Biol. Psychiatry 90 689–700

[14] Ding Z, Ouyang G, Chen H and Li X 2020 Closed-loop transcranial magnetic stimulation of real-time EEG based on the AR mode method Biomed. Phys. Eng. Express 6 035010

[15] Esposito R, Bortoletto M and Miniussi C 2020 Integrating TMS, EEG, and MRI as an approach for studying brain connectivity The Neuroscientist 26 471–86

[16] Zhang B, Liu J, Bao T, Wilson G, Park J, Zhao B and Kong J 2020 Locations for noninvasive brain stimulation in treating depressive disorders: A combination of meta-analysis and resting-state functional connectivity analysis Aust. N. Z. J. Psychiatry 54 582–90

[17] Gomez L J, Dannhauer M, Koponen L M and Peterchev A V 2020 Conditions for numerically accurate TMS electric field simulation Brain Stimulat. 13 157–66

[18] Gomez L J, Dannhauer M and Peterchev A V 2021 Fast computational optimization of TMS coil placement for individualized electric field targeting Neuroimage 228 117696

[19] Opitz A, Fox M D, Craddock R C, Colcombe S and Milham M P 2016 An integrated framework for targeting functional networks via transcranial magnetic stimulation Neuroimage 127 86–96

[20] Thielscher A, Antunes A and Saturnino G B 2015 Field modeling for transcranial magnetic stimulation: a useful tool to understand the physiological effects of TMS? 2015 37th annual international conference of the IEEE engineering in medicine and biology society (EMBC) (IEEE) pp 222–5

[21] Brainsight D T 2020 Brainsight (Montreal, Canada: Rogue Research Inc.)

[22] Brasil-Neto J P, Cohen L G, Panizza M, Nilsson J, Roth B J and Hallett M 1992 Optimal focal transcranial magnetic activation of the human motor cortex: effects of coil orientation, shape of the induced current pulse, and stimulus intensity. J. Clin. Neurophysiol. Off. Publ. Am. Electroencephalogr. Soc. 9 132–6

[23] Balslev D, Braet W, McAllister C and Miall R C 2007 Inter-individual variability in optimal current direction for transcranial magnetic stimulation of the motor cortex J. Neurosci. Methods 162 309–13

[24] MathWorks 2018 MATLAB (Natick, Massachusetts, USA: The MathWorks Inc.)

[25] Bukhari-Parlakturk N, Fei M, Voyvodic J and Michael A M 2021 Data Driven Exploration of Network Connectivity in task-fMRI of Focal Hand Dystonia medRxiv https://doi.org/10.1101/2021.05.14.21257239

[26] Voyvodic J T 1999 Real-time fMRI paradigm control, physiology, and behavior combined with near real-time statistical analysis Neuroimage 10 91–106

[27] Esteban O, Markiewicz C J, Blair R W, Moodie C A, Isik A I, Erramuzpe A, Kent J D, Goncalves M, DuPre E and Snyder M 2019 fMRIPrep: a robust preprocessing pipeline for functional MRI Nat. Methods 16 111–6

[28] Jenkinson M, Beckmann C F, Behrens T E, Woolrich M W and Smith S M 2012 Fsl Neuroimage 62 782–90

[29] Desikan R S, Ségonne F, Fischl B, Quinn B T, Dickerson B C, Blacker D, Buckner R L, Dale A M, Maguire R P and Hyman B T 2006 An automated labeling system for subdividing the human cerebral cortex on MRI scans into gyral based regions of interest Neuroimage 31 968–80

[30] CIBC 2015 Seg3D: Volumetric Image Segmentation and Visualization. Scientific Computing and Imaging Institute (SCI)

[31] Corthout E, Barker A and Cowey A 2001 Transcranial magnetic stimulation Exp. Brain Res. 141 128–32

[32] Kammer T, Beck S, Thielscher A, Laubis-Herrmann U and Topka H 2001 Motor thresholds in humans: a transcranial magnetic stimulation study comparing different pulse waveforms, current directions and stimulator types Clin. Neurophysiol. 112 250–8

[33] Bechthold, B. 2016 Violin Plots for Matlab, https://github.com/bastibe/Violinplot-Matlab

[34] Caulfield K A, Li X and George M S 2021 Four electric field modeling methods of Dosing Prefrontal Transcranial Magnetic Stimulation (TMS): Introducing APEX MT dosimetry Brain Stimul. Basic Transl. Clin. Res. Neuromodulation 14 1032–4

[35] Caulfield K A, Li X and George M S 2021 A reexamination of motor and prefrontal TMS in tobacco use disorder: Time for personalized dosing based on electric field modeling? Clin. Neurophysiol. 132 2199–207

[36] Beynel L, Dannhauer M, Plamer H, Hilbig S A, Crowell C A, Wang J E-H, Micheal A M, Wood E A, Luber B, Lisanby S H, Peterchev A V, Cabeza R, Davis S W and Appelbaum L G 2021 Network-based rTMS to modulate working memory: the difficult choice of effective parameters for online interventions. Brain and Behavior 11 e2361

[37] Awiszus F and Borckardt J 2011 TMS Motor Threshold Assessment Tool (MTAT 2.0)

[38] Smith J E and Peterchev A V 2018 Electric field measurement of two commercial active/sham coils for transcranial magnetic stimulation J. Neural Eng. 15 054001

[39] FDA 2020 Neuronal Navigator (New York City, USA)

[40] Ruohonen J and Karhu J 2010 Navigated transcranial magnetic stimulation Neurophysiol. Clin. Neurophysiol. 40 7–17

[41] Goetz S M, Kozyrkov I C, Luber B, Lisanby S H, Murphy D L, Grill W M and Peterchev A V 2019 Accuracy of robotic coil positioning during transcranial magnetic stimulation J. Neural Eng. 16 054003

[42] Schönfeldt-Lecuona C, Thielscher A, Freudenmann R W, Kron M, Spitzer M and Herwig U 2005 Accuracy of stereotaxic positioning of transcranial magnetic stimulation Brain Topogr. 17 253–9

[43] Goetz S M and Kammer T 2021 Neuronavigation The Oxford Handbook of Transcranial Stimulation. (Oxford University: Oxford University Press)

